# The impact of Arctic environments on human cerebral blood flow

**DOI:** 10.1101/2024.03.17.585446

**Authors:** Feifeng Liu, Tianheng Zheng, Jian Chen, Huazheng Liang, Gang Li, Junjie Hao

## Abstract

**Background:** The Arctic environment represents an extreme living condition that has significant impact on life. But research on alterations of human cerebral blood flow (CBF) in a high-latitude and low-temperature environment in the Arctic is still lacking.

**Methods:** Members of the 12th Chinese National Arctic Research Expedition team who took the icebreaker R/V *Xuelong* 2 to the Arctic were recruited. Transcranial colour doppler (TCD) examination was performed at the beginning of the voyage (lower latitude and higher temperature, LLHT) and during the Arctic scientific expedition period (higher latitude and lower temperature, HLLT) respectively. The spectral pattern and parameters of cerebral arteries were compared.

**Results:** Among 16 healthy individuals, 13 completed the TCD examination twice. They had a significantly lower mean velocity (Vm) (63.5 cm/s HLLT vs. 68.4cm/s LLHT; *P*=0.028) and peak systolic velocity (PSV) (94.6cm/s HLLT vs. 107.0cm/s LLHT; *P*=0.038) of the left middle cerebral artery (LMCA) and higher pulsatility index (PI) of the right middle cerebral artery (RMCA) (0.83 HLLT vs. 077 LLHT; *P*=0.011) in the HLLT environment compared to the LLHT one.

**Conclusions:** Changes in human CBF may occur in higher-latitude and lower-temperature environments in the Arctic.

## Introduction

Despite decades of research, the impact of polar environments on human physiology, especially the brain, is less well known. The only study which reported changes in the human brain was conducted in the Antarctic. In this study, the hippocampal volume of 8 participants was significantly decreased after 14-month stay in the Antarctic, so was the level of the brain-derived neurotrophic factor (BDNF). The decrease of BDNF was associated with the decreased hippocampal volume and the decline in cognitive performance in spatial processing and selective attention (Stroop Incongruent Task) tests [1]. The current study aimed to evaluate changes of the cerebral blood flow in high-latitude and extremely low-temperature environments in the Arctic.

## Methods

### Study population

Sixteen subjects were enrolled in the study. All subjects were members of 12th Chinese National Arctic Research Expedition team. They had completed comprehensive physical examinations before departure, and none of them had underlying or chronic diseases.

### Measurements

On July 11, 2021, the 12th Chinese National Arctic Research Expedition team boarded the icebreaker R/V *Xuelong* 2 from Shanghai, China, set off for a scientific expedition in the Arctic, and returned to Shanghai on September 27, 2021. The Navigation route map of the icebreaker was shown in **Figure 1**.

**Figure 1.**
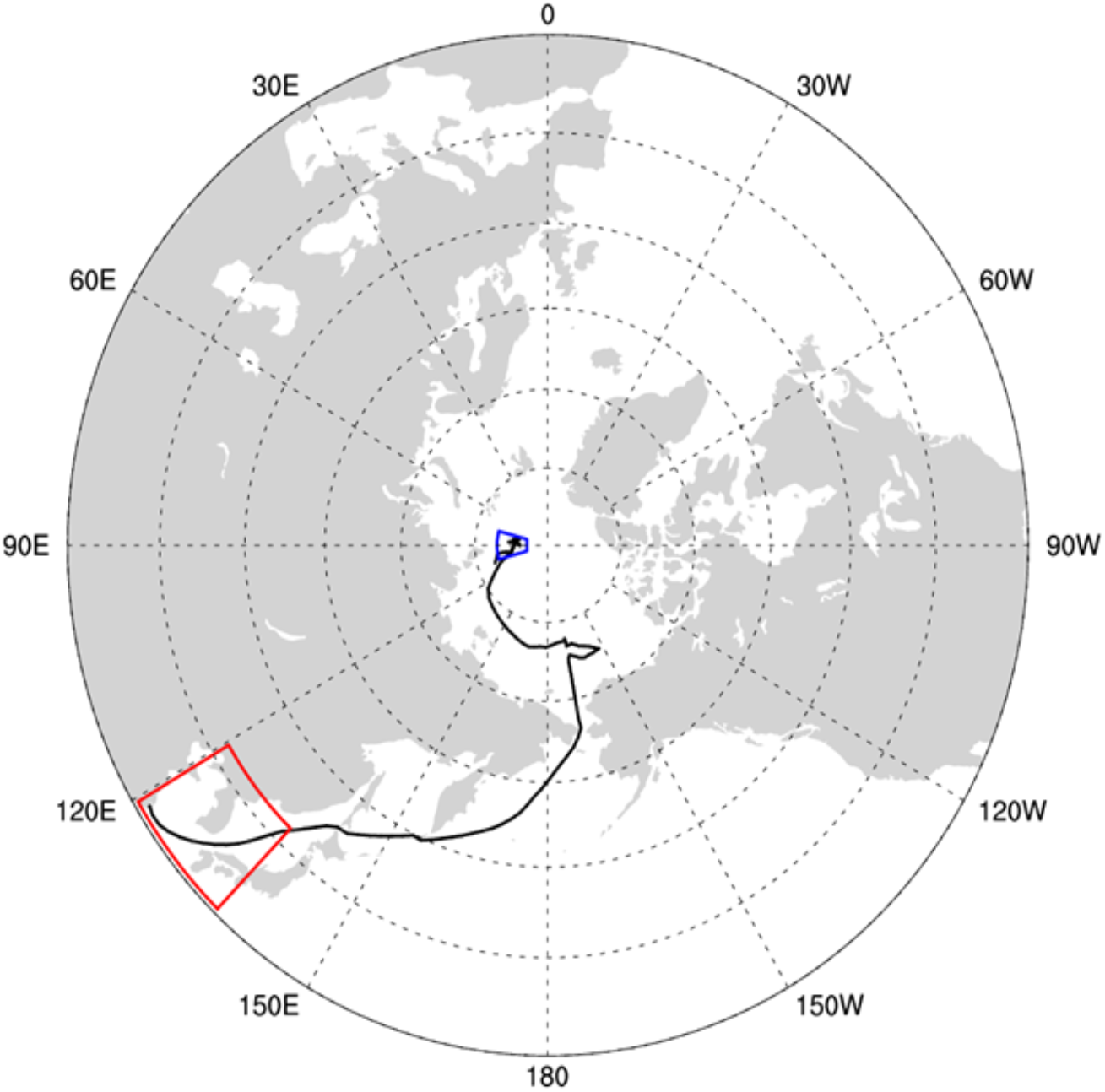
The Navigation route map of the icebreaker R/V *Xuelong* 2 during the 12th Chinese National Arctic Research Expedition. Note: The concentric circle represents the latitude line of the earth, and the outermost circle represents the 30 ° north latitude line (30 °N), which is followed by 40 °N, 50 °N, 60 °N, 70 °N, 80 °N, 90 °North Pole inward in turn; The radial dashed line represents the Earth’s longitude, E represents the east longitude, and W represents the west longitude; The red box indicates the latitude and longitude range of the icebreaker R/V *Xuelong* 2 during the first TCD inspection; The blue box indicates the latitude and longitude range of the icebreaker R/V *Xuelong* 2 during the second TCD examination; The black solid line between the red and blue boxes represents the navigation route of the icebreaker R/V *Xuelong* 2.

Transcranial colour doppler (TCD) examination was performed at the beginning of the voyage in a low-latitude and high-temperature environment (LLHT, 30°∼42° N and 120°∼140° E; Outside cabin temperature 24°C∼32°C) and during the Arctic scientific expedition period in a high-latitude and low-temperature environment (HLLT, 80°N∼90°North Pole; Outside cabin temperature - 2°C∼-6°C), respectively. The latitude and extravehicle temperature variation along the cruise of the icebreaker R/V *Xuelong* 2 during the 12th Chinese National Arctic Research Expedition was presented in **Figure 2**.

**Figure 2.**
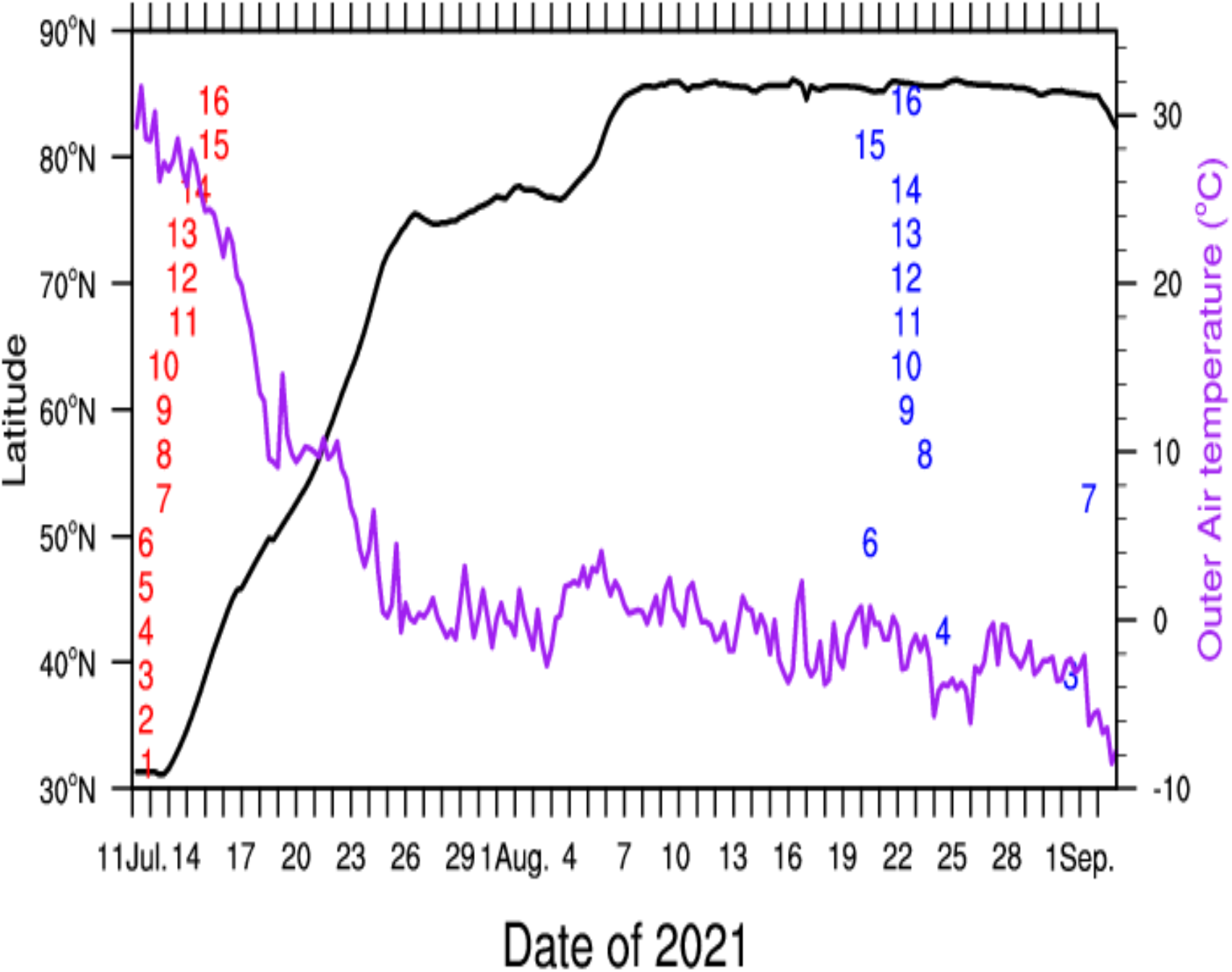
The latitude and extravehicular temperature variation along the cruise of the icebreaker R/V Xulong 2 during the 12th Chinese National Arctic Research Expedition. Note: The left longitudinal axis represents the Earth’s latitude scale, N represents the North latitude, and the right longitudinal axis represents the temperature outside the cabin in degrees Celsius (°C); The black solid line represents latitude north (°N), and the purple solid line represents the temperature outside the cabin (degrees Celsius,°C); The red Arabic numerals represent the different subjects who received TCD for the first time, and the blue Arabic numerals represent those who accepted the second TCD examination.

Cerebral blood flow parameters including the mean flow velocity (Vm), peak systolic velocity (PSV), end-diastolic velocity (EDV), pulsatility index (PI), and the resistance index (RI) were acquired using a transcranial and peripheral vascular Doppler diagnostic system (Sonara, Natus Neurology Incorporate, Ireland). Vm, PSV, and EDV values were measured from uniform blood flow patterns over 7 to 8 cardiac cycles with clear signals. PI (PI=PSV-EDV/Vm) and RI (RI=PSV-EDV/PSV) were automatically calculated.

Temporal windows were used to explore the blood flow of bilateral middle cerebral arteries (MCA), anterior cerebral arteries (ACA), and posterior cerebral arteries (PCA). Occipital windows were used to detect the blood flow of bilateral vertebral and basilar arteries.

The study was approved by the institutional ethics committee of Shanghai East Hospital, and written informed consent was obtained from each participant.

### Statistical analysis

Quantitative data were described as mean ± standard deviation (SD) if they were normally distributed or the median, i.e., interquartile range (IQR) if not normally distributed, and analyzed using paired sample t-test or Wilcoxon signed-rank test, respectively. A *p value* of < 0.05 was considered significant. All analytic procedures were conducted in SPSS version 21.0.

## Results

We initially identified 16 healthy subjects for this study. Among them, three individuals were excluded because of their refusal to take the second TCD examination. Therefore, a total of 13 subjects (all males and right-handed) with a mean age of 33 years (ranging from 22 to 48 years old) were included in the final analysis. The date when different individuals received TCD examination, the latitude of the earth where the icebreaker was located, and the temperature outside the cabin were summarized in **Figure 2**.

Parameters of cerebral blood flow for each major intracranial blood vessel were compared between the two examinations. Individuals in the HLLT environment had a significantly lower Vm (63.5 cm/s vs. 68.4 cm/s; P=0.028) and PSV (94.6 cm/s vs. 107.0 cm/s; P=0.038) of the LMCA than in the LLHT environment. However, the Vm, PSV, and EDV of the RMCA increased slightly in the HLLT environment, but these changes did not reach statistical significance. The PI of the RMCA in the HLLT environment was higher (0.83 vs. 077 LLHT; *P*=0.011) compared to that in the LLHT environment. No significant difference or trend was observed in parameters of other cerebral arteries (**Table 1**).

**Table 1.** Comparisons of TCD parameters between two different environments.

| Blood vessels | Parameters | First* (n=13) | Second† (n=13) | Difference‡ (n=13) | P value |
| --- | --- | --- | --- | --- | --- |
| LMCA | Vm, cm/s | 68.4(65.1-83.35) | 63.5(56.75-78.05) | 5.3(0.8-12.0) | 0.028 |
|  | PSV, cm/s | 107.0(83.6-107.0) | 94.6(81.55-117.5) | 9.0(0.6-16.1) | 0.038 |
|  | EDV, cm/s | 53.45±13.51 | 48.83±11.45 | 4.62±9.71 | 0.112 |
|  | PI | 0.80±0.19 | 0.76±0.12 | 0.03±0.16 | 0.461 |
|  | RI | 0.51±0.08 | 0.50±0.05 | 0.01±0.07 | 0.578 |
| RMCA | Vm, cm/s | 59.18±13.31 | 62.85±13.10 | -3.66±15.08 | 0.399 |
|  | PSV, cm/s | 90.28±20.57 | 97.27±20.56 | -6.99±23.78 | 0.310 |
|  | EDV, cm/s | 43.98±13.30 | 45.79±9.72 | -1.82±10.94 | 0.561 |
|  | PI | 0.77±0.11 | 0.83±0.09 | -0.06±0.07 | 0.011 |
|  | RI | 0.50±0.05 | 0.52±0.04 | -0.02±0.04 | 0.119 |
| LACA | Vm, cm/s | 45.88±15.61 | 42.11±18.72 | 3.78±18.43 | 0.474 |
|  | PSV, cm/s | 71.45±22.39 | 62.84±26.00 | 8.62±22.14 | 0.186 |
|  | EDV, cm/s | 33.25±12.55 | 29.95±15.48 | 3.30±13.47 | 0.395 |
|  | PI | 0.86±0.20 | 0.92±0.20 | -0.07±0.13 | 0.093 |
|  | RI | 0.51(0.495-0.59) | 0.6(0.505-0.65) | -0.03(-0.085-0.01) | 0.052 |
| RACA | Vm, cm/s | 44.87±16.00 | 43.83±13.00 | 1.04±18.11 | 0.840 |
|  | PSV, cm/s | 69.05±22.03 | 70.08±19.89 | -1.03±26.31 | 0.890 |
|  | EDV, cm/s | 33.00±13.10 | 31.55±9.47 | 1.45±14.31 | 0.721 |
|  | PI | 0.82±0.12 | 0.90±0.15 | -0.07±0.19 | 0.194 |
|  | RI | 0.53±0.05 | 0.55±0.06 | -0.02±0.07 | 0.266 |
| LPCA | Vm, cm/s | 28.75±11.76 | 27.19±7.36 | 1.55±6.59 | 0.411 |
|  | PSV, cm/s | 43.08±16.21 | 41.63±10.30 | 1.45±9.29 | 0.583 |
|  | EDV, cm/s | 21.85±9.69 | 20.11±6.04 | 1.74±5.64 | 0.288 |
|  | PI | 0.76±0.14 | 0.80±0.09 | -0.04±0.16 | 0.385 |
|  | RI | 0.50±0.06 | 0.52±0.04 | -0.02±0.07 | 0.297 |
| RPCA | Vm, cm/s | 27.16±7.00 | 22.54±3.77 | 4.62±8.14 | 0.063 |
|  | PSV, cm/s | 42.2(31.95-47.75) | 36.1(32.0-40.35) | 2.9(-4.3-16.2) | 0.289 |
|  | EDV, cm/s | 20.12±5.58 | 16.22±3.06 | 3.91±6.56 | 0.053 |
|  | PI | 0.79±0.19 | 0.87±0.16 | -0.87±0.24 | 0.215 |
|  | RI | 0.51±0.08 | 0.55±0.06 | -0.04±0.09 | 0.156 |
| LVA | Vm, cm/s | 29.63±6.57 | 34.06±12.18 | -4.43±12.26 | 0.217 |
|  | PSV, cm/s | 46.15±10.65 | 52.68±16.67 | -6.53±15.91 | 0.165 |
|  | EDV, cm/s | 21.58±4.9 | 25.04±10.31 | -3.46±10.86 | 0.273 |
|  | PI | 0.82±0.15 | 0.83±0.19 | -0.01±0.24 | 0.870 |
|  | RI | 0.52±0.06 | 0.53±0.08 | -0.003±0.10 | 0.913 |
| RVA | Vm, cm/s | 32.68±9.49 | 32.47±11.88 | 0.22±8.96 | 0.932 |
|  | PSV, cm/s | 51.31±15.39 | 50.27±17.66 | 1.04±10.45 | 0.726 |
|  | EDV, cm/s | 23.62±7.17 | 23.48±9.42 | 0.14±8.42 | 0.955 |
|  | PI | 0.85±0.18 | 0.83±0.19 | 0.02±0.19 | 0.764 |
|  | RI | 0.53±0.07 | 0.53±0.08 | 0.008±0.09 | 0.750 |
| BA | Vm, cm/s | 33.03±12.38 | 28.91±8.47 | 4.12±12.24 | 0.248 |
|  | PSV, cm/s | 51.15±20.09 | 44.85±11.44 | 6.31±18.64 | 0.246 |
|  | EDV, cm/s | 24.21±8.81 | 21.06±7.09 | 3.15±9.52 | 0.257 |
|  | PI | 0.81±0.16 | 0.84±0.15 | -0.03±0.24 | 0.618 |
|  | RI | 0.52±0.07 | 0.53±0.06 | -0.008±0.09 | 0.645 |

Data are presented as mean±SD, or median (IQR). SD indicates standard deviation; IQR, interquartile range; TCD, transcranial Doppler; LMCA, left middle cerebral artery; RMCA, right middle cerebral artery; LACA, left anterior cerebral artery; RACA, right anterior cerebral artery; LPCA, left posterior cerebral artery; RPCA, right posterior cerebral artery; LVA, left vertebral artery; RVA, right vertebral artery; BA, basilar artery; Vm, mean velocity; PSV, peak systolic velocity; EDV, end-diastolic velocity; PI, pulsatility index; RI, resistance index.
\* TCD was performed at lower latitudes and higher temperatures.
† TCD was performed at higher latitudes and lower temperatures.
‡ Difference= first – second.

## Discussion

In this observational TCD study of healthy subjects in different geographical environments, we have shown that the mean and peak systolic velocities of the LMCA were decreased and the pulsatility index of the RMCA was increased in higher-latitude and lower-temperature environments in the Arctic.

To our knowledge, this is the first investigation on the effect of terrestrial latitude and ambient temperature on changes of the velocity of the cerebral blood flow in humans. There is little research on the impact of environmental factors on human brain functions in polar regions (including both the North and the South Poles). Only German scholars have explored Brain Changes in Response to Long Arctic Expeditions in 2019^[1]^. They assessed changes in the volume of hippocampal subfields, whole-brain grey matter, and brain-derived neurotrophic factors in venous blood samples. They found a decrease in the hippocampal volume of the dentate gyrus, the volume of the grey matter in the right dorsolateral prefrontal cortex, the left orbitofrontal cortex, and the left parahippocampal gyrus, and a decreased in the concentration of the brain-derived neurotrophic factor after a long duration of overwintering in the Antarctica.

There are some notable limitations in this study. Only thirteen participants were included in our study. The study duration is relatively short, with a time interval between two TCD examinations only over one month. It took study subjects one month to arrive at the Arctic region where they stayed for another month and a half, which may be not sufficient to cause significant changes in humans. In addition, due to limited conditions, the research subjects are only adult males and they can not fully reflect the physiological changes in the human of different genders. Finally, only TCD was used to evaluate the cerebral blood flow, with few other examination items, which renders us unable to fully characterize changes in physiological functions of the human brain. Therefore, our data should be interpreted with caution as no data are available to explain the underlying mechanism. It is necessary to develop long-term research, including both male and female studies, using more tools in the future to explore changes in the human in polar environments, if conditions permit.

In summary, this observational study identified that human cerebral blood flow may decrease in high-latitude and low-temperature environments in the Arctic.

## Contributors

LFF and ZTH conceived and designed the study, wrote the first draft of the manuscript and analyzed the data. HJJ is one of the members of the Chinese 12th Arctic Expedition team responsible for the study design, data collection. LG obtained the funding, supervised the study and finalized the manuscript. HL revised the manuscript. All authors revised and approved the final manuscript.

## Data availability

Non-published anonymized data can be shared with qualified investigators upon reasonable request to the corresponding author.

## Acknowledgments

We thank all the members of the Chinese 12th Arctic Expedition team for their continuous support of this study.

